# Toxicogenomic Assessment of In Vitro Macrophages Exposed to Profibrotic Challenge Reveals a Sustained Transcriptomic Immune Signature

**DOI:** 10.1101/2024.06.19.599702

**Authors:** Jack Morikka, Antonio Federico, Lena Möbus, Simo Inkala, Alisa Pavel, Saara Sani, Maaret Vaani, Sanna Peltola, Angela Serra, Dario Greco

## Abstract

Immune signalling is a crucial component in the progression of fibrosis. However, approaches for the safety assessment of potentially profibrotic substances, providing information on mechanistic immune responses, are underdeveloped. This study utilises a comprehensive analysis of RNA sequencing data from macrophages exposed in vitro to multiple sublethal concentrations of the profibrotic agent bleomycin, over multiple timepoints. Using a toxicogenomic framework, we performed dose-dependent analysis to filter genes truly altered by bleomycin exposure from noise and identified a subset of immune genes with a sustained dose-dependent and differential expression response to profibrotic challenge. We performed an immunoassay and revealed cytokines and proteinases responding to bleomycin exposure that closely correlate to transcriptomic alterations, underscoring the integration between transcriptional immune response and external immune signalling activity. This study not only increases our understanding of the immunological mechanisms of fibrosis, but also offers an innovative framework for the toxicological evaluation of substances with potential fibrogenic effects on macrophage signalling. Our work brings a new immunotoxicogenomic direction for hazard assessment of fibrotic compounds, through implementation of a time and resource efficient in vitro methodology.

## 1. Introduction

Pulmonary fibrosis (PF), originating from interstitial lung diseases, encompasses a range of underlying aetiologies. Occupational exposures to asbestos, silica, metal dusts, and nanoparticles such as multi-walled carbon nanotubes are well-recognized causal events (1–4). Viral infections, for example SARS-CoV-2, have emerged as potent triggers for fibrotic alterations to the lung (5,6). Drug-induced fibrosis, from agents such as bleomycin, represent a notable iatrogenic cause (7). There are also cases of idiopathic PF, aggravated by lifestyle factors such as smoking, but where the causative factors remain elusive (8,9). Whilst the aetiology of PF is multifaceted, each initiating event shares a common subsequent step in the progression to fibrosis, whereby immune cells are recruited to clear the profibrotic insults and to resolve damage to the epithelial cell layer (10). These immune cells externally signal for the recruitment of fibroblasts, which are the principal effector cells driving fibrosis (11). An overproduction of extracellular matrix components (ECM), particularly collagen, fibronectin and hyaluronic acid, ultimately leads to scarring and the impairment of gas exchange and diminished lung function (12,13). Despite knowledge of a common immune response to varied profibrotic exposures in PF, efforts to develop a means to rigorously assess the mechanistic immunotoxic potential of fibrogenic substances are lacking.

The most prominent immune cells, both in number and activity in the lung, are macrophages (10,14). Lung macrophages are broadly categorised into alveolar and interstitial macrophages (15). Recruited monocytes that differentiate into macrophages at the site of the lung have been shown in murine models to be important in driving pulmonary fibrosis, whereas alveolar macrophages had no effect (16). Similarly, CCR2 full knock out mice, missing a receptor required for monocyte infiltration into the lung, are also protected from lung fibrosis (17,18). One of the primary forms of macrophage communication is via cytokine signalling. TGFB is the most studied cytokine in the context of pulmonary fibrosis. Acting through SMAD signalling and the JNK/MAPK pathways, it promotes the production of fibronectin, proteoglycans, and collagen and inhibits the degradation of ECM by matrix metalloproteinases (19,20). The interleukins IL-1B, IL-17A and IL-18 are well known pro-inflammatory interleukins elevated in both IPF patients and murine models of pulmonary fibrosis (21,22). However, whilst the importance of macrophages and macrophage cytokine signalling is well understood in the progression of PF, a comprehensive understanding of the mechanistic response of macrophages to profibrotic challenge is understudied. The ability of profibrotic compounds to alter the immune activity of macrophages at a mechanistic level is yet to be fully explored. The traditional understanding of M1 and M2 macrophage polarization, whereby macrophages take on pro-inflammatory and anti-inflammatory states (23,24), is overly simple in the context of lung fibrosis, as shown by the recent availability of single cell data, where distinct populations of macrophages are observed in the fibrotic niche of lung tissue from IPF patients (25–28). That said, these single cell studies are also limited to only observing macrophage transcriptomes at the end stages of PF and not informing on acute, early-stage responses by macrophages when they have just arrived at the lung to clear the initial profibrotic insult. It is this early interaction with profibrotic agents that activates the macrophages into an altered state capable of driving fibrosis, and which should be the focus of attempts to scrutinise the alteration of macrophage activity in response to profibrotic substances. Thus, gaining a mechanistic understanding of the early-stage modulation of immune signalling by profibrotic agents in the context of lung fibrosis is paramount, particularly for the field of chemical safety assessment where it is the initial cellular responses to a substance that are of interest and determine the substance’s toxic profile. Greater mechanistic understanding of these early stages necessitates the development of time-efficient testing methodologies that accurately encompass the initial phase of disease progression, ensuring that the potential profibrotic agents are evaluated within a relevant early-stage of pathological development.

Bleomycin is a cytotoxic glycopeptide derived from the bacterium *Streptomyces verticillus* and used as an antineoplastic in the treatment of Hodgkin’s Lymphoma, testicular cancer and in other chemotherapeutic regimes (29–31). Bleomycin’s cytotoxic activity is onset by its ability to cause single- and double-stranded DNA breaks. Due to the high oxygen environment in the lung enabling effective oxidative damage by bleomycin to DNA, and due to the lack of bleomycin hydrolase to enzymatically degrade bleomycin in lung cells, treatment with bleomycin often leads to its most serious side effect of pulmonary toxicity and pathological levels of fibrosis in the lung (7,32,33). As a well-known profibrotic agent, bleomycin is commonly used in rodent studies to model the onset of PF with its associated hallmarks of epithelial damage, inflammation and altered immune signalling, epithelial– mesenchymal transition (EMT), and myofibroblast activation all consistently onset by tracheal instillation of bleomycin (34–36).

To model the early effects of a direct profibrotic insult on macrophage activity and immune signalling, we exposed differentiated THP-1 macrophages, a model of monocyte derived macrophages (37,38), to bleomycin at multiple sublethal concentrations over multiple timepoints, generating what can be considered fibrotically activated macrophages (FAMs). Employing an approach used in toxicogenomics, we performed dose-dependent analysis to filter genes truly altered by bleomycin exposure, from noise (39,40). Our results indicated a cohort of 108 differentially and dose-dependently expressed immune genes showing consistent expression changes over 72 H of bleomycin exposure. We also measured a panel of cytokines and proteinases, known to be important in the progression of fibrosis. We correlated expression the implicated cytokine-proteinases to the expression of the differentially and dose-dependently regulated genes, highlighting a close link between immune gene regulation and cytokine and proteinase secretion by the FAMs. The panel of immune genes uncovered using this framework represents an early immune signature caused by profibrotic exposure. Accordingly, the methodology described herein—subjecting macrophages to sublethal concentrations of profibrotic compounds, followed by RNA sequencing with dose-dependent analysis and subsequent correlation of cytokine and proteinase profiles across multiple timepoints—embodies an efficient and resource-conservative approach for the safety assessment of substances with potential profibrotic and immunotoxic effects.

## 2. Materials and Methods

### Cell culture

THP-1 cells (ATCC TIB-202, USA) were cultured in RPMI-1640 (Gibco, USA) supplemented with 10% FBS (Gibco, USA) (culture media). Cells were cultured in 75 cm^2 flasks at 37 °C with a humidified atmosphere of 5% CO2, at a density < 1 × 106 cells/mL. Cells were plated and differentiated in 12 well plates at a density of 0.445 x 10^6 cells/mL (148333 cells/cm2), in culture media supplemented with 50 nM of phorbol 12-myristate 13-acetate (PMA) (Sigma-Aldrich, USA) for 48 H prior to bleomycin exposure.

### Bleomycin exposure

THP-1 cells were exposed to 0-100 µg/mL of bleomycin ready-made solution (Sigma-Aldrich, #B7216) in 1 mL of THP-1 culture media, using water as the vehicle at 1.8% v/v concentration for all conditions including 0 µg/mL control, with continuous exposure for either 24, 48 or 72 hours. Exposure was randomised on 12 well plates using the R package Well Plate Maker (41).

### WST-1 viability assay

For the viability assay there were 4 samples for each concentration of bleomycin (0,20,40,60,80, and 100 µg/mL) at each time point (24, 48 or 72 hours). A WST-1 assay was used to measure cell viability. Briefly, 100 µL of cell proliferation reagent WST-1 (Roche, #11 644 807 001) was added to each well. Cells were left to incubate with WST-1 for 2 H in a 37 °C, 5% CO2 incubator. Absorbance at 450 nm was measured with a Spark microplate reader (Tecan). Full results can be found in **Supplementary File S1**.

### Procartaplex Immunoassay

Supernatant was collected for cytokine and proteinase profiling using a custom 22-plex Procartaplex assay (ThermoFisher, #PPX-22-MXRWGMP, custom kit) for the following targets: CXCL11, FGF-2, IFN gamma, IL-1 alpha, IL-1 beta, IL-10, IL-13, IL-17A, IL-18, IL-6, IL-8, IP-10, MCP-1, MIG, MIP-1 alpha, MMP-1, MMP-7, MMP-9, PDGF-BB, SDF-1 alpha, TNF alpha, VEGF-R2. Supernatant was collected from the same samples used for RNA extraction for RNA sequencing, and frozen at −80 °C. Supernatant samples were thawed on ice, vortexed, and centrifuged at 10,000 x g for 5 minutes prior to analysis. Standards, blanks, and samples were prepared according to manufacturer’s instructions and were dispensed into designated wells in a random layout created using the R package Well Plate Maker (41). Results were measured using a Bio-plex 200 system (Bio-Rad). The following targets were undetected in the range of the standard curves and therefore could not be used for further analysis: CXCL11, IL-13, IL-8, MIP-1 alpha, VEGF-R2. For each detected target, background (blank) fluorescence was subtracted and the remaining fluorescence was extrapolated to the relevant standard curve to derive concentration measurements in pg/mL. Full immunoassay results can be found in

### Supplementary File S1. RNA extraction

For both RNA sequencing and the independent repeat experiment for qPCR validation, there were 4 samples for each concentration of bleomycin (0,20,40,60,80, and 100 µg/mL) at each time point (24, 48 or 72 hours). Media was removed and cells were washed briefly with 500 µL of cold PBS. 350 µL of lysis buffer from the QIAGEN RNeasy mini kit (Qiagen, #74104) was added to each well to lyse the cells. Total RNA was then extracted from these samples using the QIAGEN RNeasy mini kit (Qiagen, #74104) following manufacturer’s instructions. DNase treatment was performed using DNase1 RNaseFree (ThermoFisher, #EN0521) according to the manufacturer’s instructions.

### RNA sequencing

RNA sequencing was performed by the company Novogene. Quantity and quality of the RNA samples were assessed with their in-house quality checks as follows: preliminary quality control was performed on 1% agarose gel electrophoresis to test RNA degradation and potential contamination, sample purity and preliminary quantitation and RNA integrity were measured using Bioanalyzer 2100 (Agilent Technologies, USA). For library preparation, the Novogene NGS RNA Library Prep Set (PT042) was used. The mRNA present in the total RNA sample was isolated with magnetic beads of oligos d(T)25 using polyA-tailed mRNA enrichment. Subsequently, mRNA was randomly fragmented and cDNA synthesis using random hexamers and reverse transcription was performed. Once first chain synthesis was finished, the second chain is synthesised with the addition of an Illumina buffer (non-directional library preparation). Together with the presence of dNTPs, RNase H and polymerase I from E. Coli, the second chain was obtained by Nick translation. Resulting products then underwent purification, end-repair, A-tailing and adapter ligation. Fragments of the appropriate size were enriched by PCR, where indexed P5 and P7 primers (Illumina) were introduced, and final products are purified. The library was checked with Qubit 2.0 and real-time PCR for quantification and bioanalyzer Agilent 2100 for size distribution detection. Quantified libraries were pooled and sequenced on the Illumina Novaseq X platform, according to effective library concentrations and data amounts using the paired-end 150 strategy (PE150).

### RNA Sequencing Analysis

Raw sequencing reads were subjected to initial quality assessment using FastQC v0.11.7 (42), followed by the trimming of Illumina adapters and the removal of low-quality bases using TrimGalore v0.4.4_dev (43). Subsequent quality control was reassessed using FastQC v0.11.7. The processed reads were aligned to the human reference genome (GRCh38) using HISAT2 v2.1.0 (44). Post-alignment, BAM file indexing was performed and uniquely mapped reads were generated using SAMtools v1.8-27-g0896262 (45). Gene-level read count matrices were generated using the featureCounts function from the package Rsubread v1.34.6 (46). Low expressed genes were filtered out by applying a proportion test, as implemented in the R package NOIseq v2.28.0 (47). The raw sequencing data and pre-processed counts matrices are accessible at: https://doi.org/10.5281/zenodo.11371243. For exploratory data analysis, including principal component analysis (PCA) to visualize data structure and upset plotting to explore intersections of differentially expressed genes across conditions, reads were normalized using the variance stabilizing transformation (VST) method from the R package DESeq2 v1.34.0 (48). Differential expression analysis was performed using DESeq2 v1.34.0, modelling gene expression as a function of bleomycin concentration contrasted to concentration 0 µg/mL. Significantly differentially expressed genes were identified at an adjusted p-value (FDR) ≤ 0.01 and an absolute log2 fold change ≥ 0.58 (equivalent to a fold change of ≥ 1.5). Enrichment analysis of significantly differentially expressed genes to analyse altered biological pathways and processes was performed using the clusterProfiler package (49).

### Dose-Dependent Expression Analysis Using BMDx

Dose-dependent gene expression in THP-1 macrophages exposed to bleomycin was assessed using BMDx software (39). Counts matrices for each time point (24, 48, and 72 hours) was transformed using DESeq2 Variance Stabilizing Transformation (VST) and used as input. Selection of dose-dependent genes was based on an R-squared (R^2) threshold of 0.6 to ensure a robust dose-response relationship. Statistical models, specifically exponential (exp2), hill, linear, polynomial (poly2), and power models, were fitted to the VST-transformed RNA expression data from RNA sequencing to determine dose-dependent expression, and the model with the lowest Akaike Information Criterion (AIC) for each gene was selected. The BMD and BMDL values were computed from these models were utilized to evaluate the fidelity of model fittings at each timepoint. The same procedure, with the same parameters and models fitted (with the addition of Log-Logistic Model (llog2), Michaelis-Menten Model (mm2) and Weibull (weibul12)), was used to perform dose-dependent analysis on the log2FC expression changes of the procartaplex immunoassay cytokine and proteinase targets that passed an ANOVA filtering with an adjusted p value <0.05 (**Supplementary File S1**).

### Correlation Analysis of Gene and Cytokine-Proteinase Expression

To examine the relationship between gene expression and cytokine-proteinase secretion in response to bleomycin, a Spearman correlation analysis was performed (**Supplementary File S1**). Log2 fold change (log2FC) data for both gene expression and cytokine-proteinase secretion were used. The gene expression data encompassed differentially expressed and dose-dependent (DE∩DD) genes in THP-1 macrophages exposed to bleomycin (0-100 µg/mL). The cytokine-proteinase data included log2FC values for eight identified proteins that were significantly differentially secreted (ANOVA p <0.05, **Figure S4**, **Supplementary File S1**). The correlation at each timepoint (24, 48, and 72 hours) was analysed separately. Correlations were calculated using the Spearman’s rho, filtering for strong associations (|ρ| > 0.8) and adjusted for multiple testing using the false discovery rate (FDR, p < 0.05). Significant correlations were further filtered for genes involved in the immune system process (GO:0002376), ensuring relevance to immunological responses in our analysis. Full correlation results can be found in **Supplementary File S1**.

### BiomarkHD qPCR validation

qPCR was performed for bleomycin concentrations 0,20,80, and 100 µg/mL. Synthesis of cDNA, from 277 ng of DNase treated RNA for each sample, was performed using the high-capacity cDNA reverse transcription kit (Thermo Fisher Scientific, #4368813), according to manufacturer’s instructions. Expression levels of target genes were determined by high-throughput multiplex qRT-PCR using the BiomarkHD and IFC Controller MX system (Standard Biotools). Primers used were D3 deltagene assays designed and ordered with Standard Biotools (**See Supplementary File S1**). Fluidigm Preamp Master Mix (Fluidigm, #100-5581), Exonuclease 1 (ThermoFisher, #EN0581), SsoFast EvaGreen Supermix with Low ROX (Bio-Rad, #1725211) and a Deltagene chemistry; EG=Eva Green kit (AH Diagnostics, #BMK-M10-48.48-EG) were used with a 48.48 integrated fluidic circuit (IFC) chip, according to the BiomarkHD manufacturer’s instructions. Preamplified cDNA was diluted 5-fold in TE buffer [1mM EDTA] (Invitrogen, #AM9849) and the BiomarkHD thermal protocol GE Fast 48×48 PCR+Melt v2 was used for the final run. Specificity of single site amplification was confirmed by performing a melt curve analysis. GeNorm analysis (50) of 6 reference genes (PUM1, RPL37A, SNW1, IPO8, GAPDH, ACTB) was performed using R packages ReadqPCR v1.40.0 and NormqPCR v1.40.0 (51) to assess and select the most stable reference genes for relative expression analysis. Fold change (FC) values from RT-qPCR data were calculated using the comparative CT(2−(ddCt)) method (52). The FC values were log2 transformed (log2(FC)). For each gene and for each concentration, an outlier detection was performed by removing all the samples with log2(FC) values above or below the 75th and 25th percentiles of the distribution. Ct values, dCt values, FC values and log2(FC) values and full results are available in **Supplementary File S1**, along with ANOVA tables and tukey HSD posthoc test results.

### Discriminant Fuzzy Pattern Analysis of DE∩DD immune genes

To estimate baseline expression of the 108 DE∩DD immune genes, single cell gene expression data from the tabula sapiens dataset (53), was queried from a previously introduced Knowledge Graph (KG) framework (54), concentrating on tissue specific macrophages at the steady state, and transformed into categorical values using the Discriminant Fuzzy Pattern (DFP) method (55). Briefly, continuous gene expression scores were classified into low, medium, or high categories by fitting Gaussian functions to each dataset and gene, estimating class probabilities. Genes were assigned class labels if the probability was ≥0.5; genes with an expression value of zero across all samples were considered non-measured. In case of genes with multiple labels, priority was given to higher labels (for example, if a gene would have both high and medium label, high would be selected). Expression data for 74 of the 108 genes was present in the tabula sapiens dataset.

### Ranking analysis of DE∩DD immune genes

The ranking of genes in **Table S1** was determined using a comprehensive scoring system for the panel of 108 differentially expressed and dose-dependent (DE∩DD) immune genes responding to bleomycin exposure. Each gene received a score based on multiple criteria: 1) correlation with cytokine-proteinase secretion, 2) identification as a cytokine or cytokine receptor, 3) Implication in PF as listed in the DisGeNET database (56), which contains information on genes associated with human diseases, 4) presence in genome-wide association studies (GWAS) data for idiopathic pulmonary fibrosis (IPF) retrieved from the GWAS catalog (57), 5) status as a target for approved drugs retrieved from the Open Targets Platform (58), 6) frequency of implication in eleven harmonised public RNA sequencing datasets comparing biopsies from IPF patients to healthy lung tissue (harmonised data available at: https://doi.org/10.5281/zenodo.10692129); briefly, RNA sequencing datasets were retrieved from the European Nucleotide Archive (ENA); differential gene expression was performed with DESeq2 v1.34.0 (48); in total, 629 samples, consisting of 360 disease samples and 269 healthy samples, were represented across the 11 datasets; and a score for **Table S1** from 0 to 1.1 in steps of 0.1 was given depending on how many datasets each gene was represented in. The scores from all 6 of these categories were finally summed to generate a rank for the prioritization of genes according to their relevance in the immunotoxic signature of macrophages exposed to bleomycin.

## 3. Results and Discussion

We selected bleomycin as our chosen profibrotic agent to develop a toxicological safety assessment framework for compounds with immune altering fibrogenic potential. The fibrosis caused by bleomycin exposure is mediated in part by the continuous recruitment and infiltration of monocyte derived macrophages (MDM) (16). Despite the known role of MDM in the progression of PF, the mechanistic response of these cells upon direct interaction with a profibrotic insult such as bleomycin, is yet to be fully elucidated. To gain further insight into how profibrotic insults might be affecting the immune activity of macrophages, and to generate a model to assess immunotoxicity of profibrotic compounds, we exposed differentiated THP-1 macrophages to multiple doses of bleomycin (0-100 µg/mL) over multiple timepoints (24 H,48 H,72 H). We then analysed the mechanistic processes and immune signature of these fibrotically activated macrophages (FAMs) by RNA sequencing and immunoassay (**Figure 1**).

**Figure 1:**
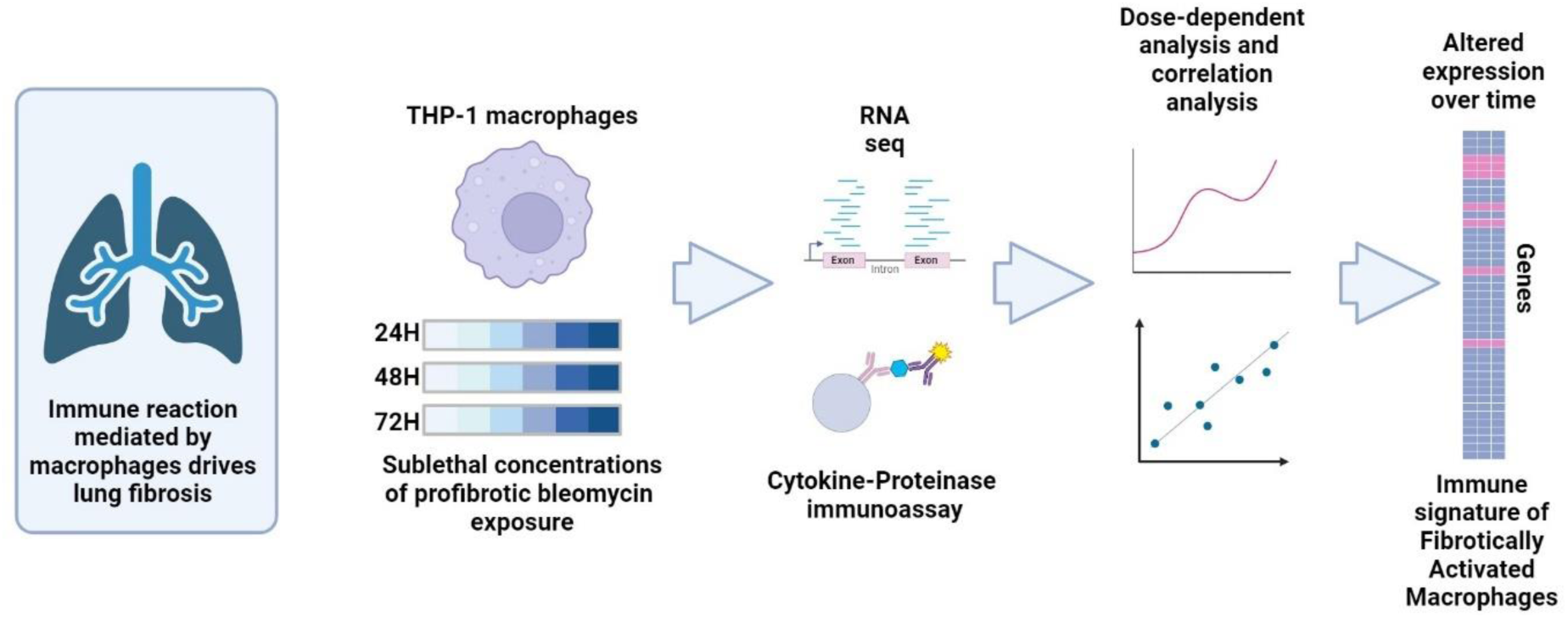
Schematic representation of the study. Lung fibrosis is driven in part by an excessive immune reaction mediated by macrophages. This study develops a framework to assess the effect of profibrotic exposure on the immune signature of fibrotically activated macrophages. THP-1 macrophage cells were exposed in-vitro to 0-100 µg/mL of bleomycin. RNA sequencing and cytokine immunoassays were performed and analysed using dose-dependent modelling. Correlation of gene expression to cytokine expression revealed genes likely to be controlling cytokine expression. Immune gene filtering of both the RNA sequencing analysis and the correlation analysis reveals a distinct immune gene expression signature in response to bleomycin, sustained across timepoints.

### 3.1 Sublethal concentrations of bleomycin exposure leads to dose-dependent shifts in macrophage gene expression

We first established a sublethal range for continuous exposure to bleomycin over a 72-hour period (**Figure S1**). We ensured that the viability at the highest concentration (100 µg/mL) did not go below 75% of that observed in the vehicle control, to prevent the masking influence of gene expression changes associated with cell death, which can obscure the underlying mechanistic insights into bleomycin exposure in downstream RNA sequencing analysis (59). We then used these sublethal concentrations and reran the same experiment, exposing THP-1 macrophages to bleomycin but this time extracting RNA at all 3 timepoints (24 H,48 H,72 H) for sequencing. PCA plots from the resultant RNA count matrices show a visible dispersion of samples across PC1, indicating a concentration-dependent separation in gene expression profiles, with each concentration significantly distinct from the control (**Figure S2A**). At 48 H, the variance explained by PC1 increases to 83%, and by 72 H, PC1 explains 89% of the variance, with a tighter cluster of low-concentration treatments indicating less variation at these levels over time (**Figure S2A**). The clear distinction between concentrations, with the highest concentrations showing the greatest displacement along PC1, indicates a pronounced gene expression response due to multi-dose exposure to bleomycin, with greater resolution of the variation within and between groups over time (**Figure S2A**). At 24 H, the number of differentially expressed (DE) genes is relatively modest at lower concentrations, but there is a linear increase in DE genes peaking at the highest concentration (100 µg/mL). This linear increase in DE genes across concentrations is also observed at 48 H and 72 H. Moreover, the total number of DE genes also increases as the length of time of continuous exposure increases, indicating a linear increase in DE genes not only across concentrations, but also across timepoints (**Figure S2B**).

Whilst differential expression analysis indicated genes that are DE at specific concentrations, many of these genes do not follow a dose-dependent pattern of expression and instead represent false-positives, or noise. Due to the redundancy and resilience of regulatory gene expression circuits, simple RNA expression analysis of experiments designed using single dose, treated vs untreated setups, misses crucial information generating both false positive (e.g. differentially expressed but only randomly at a single dose) and false negative results (e.g. differentially expressed at other exposure concentrations but not detected at the single concentration measured). Novel toxicogenomic approaches have begun to use statistical modelling to distinguish genes that follow a dose-dependent pattern of expression, thereby highlighting dose-dependent (DD) genes that can confidently be said to be perturbed by an exposure (39,40). DD gene expression indicates that the gene is under regulatory control that responds quantitatively to the level of exposure and not just a binary on/off expression. Through the detection of DD genes, we are discovering genes that are directly affected by the exposure itself and not as an indirect response to exposure resulting from the inherent redundancy and resilience of gene signalling pathways, ensuring we focus on assessing and detecting the most relevant genes in the context of a toxicological safety assessment. When applied to the count matrices from our bleomycin exposure, DD analysis discovered 1723, 2299 and 4646 DD genes at 24 H, 48 H, and 72 H respectively, fitting common statistical models and displaying consistent BMD/BMDL ratios indicating few outliers and consistent quality of model fitting (**Figure S3**). Many of these DD genes may not be DE genes in our analysis of differential expression, however, and as such, for each timepoint, we found the intersection of DE genes with DD genes (**Figure 2A**). This DE∩DD intersection represents the macrophage genes meaningfully responding to bleomycin exposures in a dose-dependent fashion.

**Figure 2:**
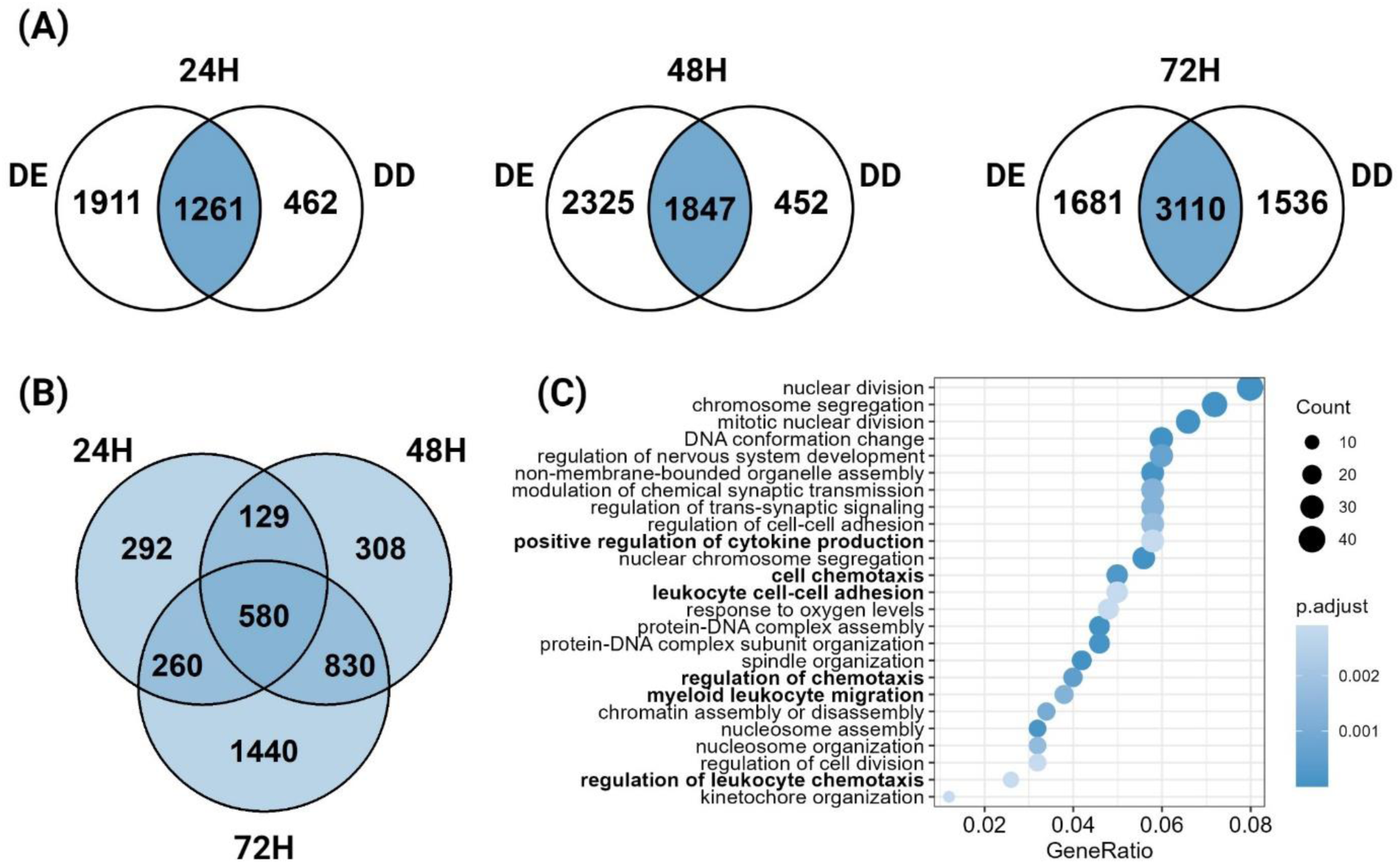
Analysis of RNA sequencing data for THP-1 macrophages exposed to 0-100 µg/mL of Bleomycin at 24 H, 48 H and 72 H. (A) The intersection (∩) of differentially expressed (DE) and dose-dependent (DD) genes responding to bleomycin at each timepoint is indicated in blue (B) A Venn diagram shows the subsection of DE∩DD genes sustained across all three timepoints. (C) Enrichment analysis of DE∩DD genes sustained across all three timepoints. Count shows the number of genes represented in each ontology. Gene ratio shows the ratio of genes in this ontology to all genes used for this enrichment (580 in total). Adjusted p-value is indicated by shade of blue. Ontologies that are immune related (GO:0002376 Immune System Process) are emboldened.

Our analysis highlighted that 580 of these DE∩DD genes were commonly dysregulated at all 3 timepoints (**Figure 2B**), representing a subset of genes that have an acutely onset and sustained response to bleomycin exposure in macrophages. Enrichment analysis of these genes primarily indicated terms involved in the internal response to the DNA-damage and cytostatic mechanism of action (MOA) of bleomycin. With biological processes including, *nuclear division, chromosome segregation and DNA conformation change* in the top 5 significant categories (**Figure 2C**). Whilst these processes would otherwise indicate ongoing cell division, the bleomycin exposed THP-1 cells are in a differentiated and non-proliferating state, thus these terms instead represent pathways involved in repairing the DNA damage caused by bleomycin. In the progression of PF however, and when assessing the fibrogenic potential of a compound (in this case bleomycin), it is the external immune activity of macrophages, such as cytokine signalling, that are of interest regarding downstream fibroblast and immune cell recruitment and activation, ultimately resulting in pulmonary toxicity. The immune categories *positive regulation of cytokine production*, *cell chemotaxis, leukocyte cell-cell adhesion*, *regulation of chemotaxis*, *myeloid leukocyte migration* and *regulation of leukocyte chemotaxis*, were all found in the top 25 significantly enriched biological processes for the 580 sustained DE∩DD genes (**Figure 2C**). Chemotaxis is an essential process in the recruitment of immune cells and fibrocytes to the site of damage in PF. Bronchoalveolar lavage fluid from patients with IPF shows increased immune cell chemotactic activity (60), with IL-8, CXCL1 and CCL18 cytokines showing potent immune cell chemoattractant activity (61). A subpopulation of lung interstitial macrophages residing next to blood vessels increase recruitment and infiltration of further monocyte derived macrophages, mediated in part by CCL3 (MIP-1α) and CCL22 chemotactic cytokine secretion, creating a cycle of increasing immune cell infiltration and fibrosis progression (62). Thus, the chemotaxis and cytokine regulation categories observed in our enrichment analysis of bleomycin responding DE∩DD genes was expected, but to assess in detail the immunotoxicity of bleomycin exposure in the context of chemical safety assessment, the immune component of the transcriptomic response deserved further exploration.

### 3.2 Toxicogenomic assessment of immunotoxicity in fibrotically activated macrophages

We focussed on the immune response of macrophages to bleomycin exposure by filtering the DE∩DD genes for immune genes only, belonging to the gene ontology GO:0002376, immune system process (**Figure 3A**). Our analysis revealed a cohort of 108 immune-specific DE∩DD genes that consistently responded to bleomycin across all three examined time points (**Figure 3B, Table S1**). This consistent response underscores a sustained immune signature characteristic of macrophages exposed to a profibrotic challenge. Further investigation into the biological processes enriched among these 108 immune genes again highlighted the significance of leukocyte chemotaxis and migration, alongside cytokine signalling, in mediating the macrophage response (**Figure 3C**). Our functional analysis also featured a pronounced enrichment in T-cell activation-related processes. Specifically, the terms *T cell activation*, *regulation of T cell activation*, and *positive regulation of T cell activation* were among the top 25 significantly enriched categories (**Figure 3C**). The exact role of T cell activity in PF remains unclear despite research showing their involvement (63). Knockout of γδ T cells in mice can increase the severity of bleomycin induced PF, decreasing the concentration of IL-6, CXCL1, and CXCL10 suggesting a role of these specific T cells in mitigating fibrosis (64). Linked to this, the entire family of BTN3A butyrophilins (BTN3A1, BTN3A2, BTN3A3) were down regulated in the 108 immune-specific DE∩DD genes at all 3 timepoints in our bleomycin exposed macrophages (**Figure 5**). Whilst these butyrophilins activate γδ T cells in an intracellular manner in response to phosphoantigens, they are also found on the surface of multiple cells types (including macrophages), regulating the activity of γδ T cells by binding their T cell receptor (65). Our FAMs are potentially down-regulating BTN3A expression to mitigate γδ T cell activity and drive fibrosis for wound repair. Our results show that FAMs likely play an important role in regulating T cell activity in response to profibrotic insult (11). In summary, by focussing in on the immune response of macrophages to bleomycin, we illustrate how toxicogenomic assessment in the form of dose-dependent and differential analysis of RNA sequencing data does not only inform on the immunotoxic alterations of significant immune processes, but can also highlight specific dysregulated biological activities that may have particular relevance for complex diseases that are known outcomes of the substances being tested, such as PF.

**Figure 3:**
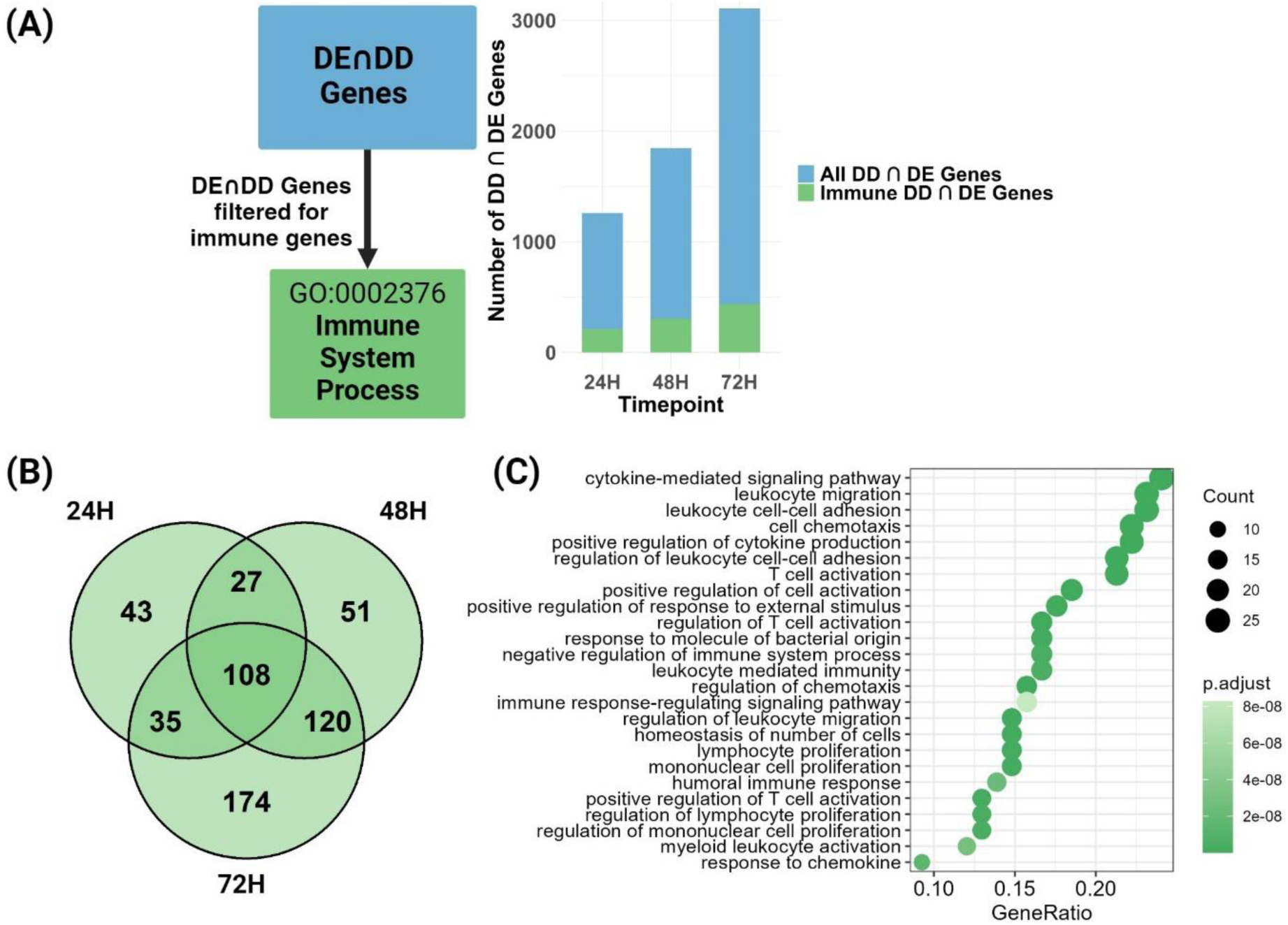
Immune gene expression in THP-1 macrophages exposed to 0-100 µg/mL of Bleomycin at 24 H, 48 H and 72 H. (A) The intersection (∩) of differentially expressed (DE) and dose-dependent (DD) genes responding to bleomycin at each timepoint (blue) is filtered for immune genes found under GO:0002376 (green). A barplot shows the proportion of immune genes (green) from total DE∩DD genes (blue) at each timepoint (B) A Venn diagram shows the subsection of immune DE∩DD genes sustained across all three timepoints. (C) Enrichment analysis of immune DE∩DD genes sustained across all three timepoints. Count shows the number of genes represented in each ontology. Gene ratio shows the ratio of genes in this ontology to all genes used for this enrichment (108 in total). Adjusted p-value is indicated by shade of green.

### 3.3 An immune gene response to profibrotic insult controls external cytokine-proteinase secretion

Macrophages orchestrate in-situ immune activity, and the recruitment of immune cells and fibroblasts, through the release of cytokines and contribute to the remodelling of the extracellular matrix (ECM) by secreting proteinases, thereby facilitating functional and structural alterations necessary for tissue repair and fibrosis. Thus, a comprehensive assessment of the profibrotic potential of test substances should encompass an analysis of cytokine activity. We selected a panel comprising 14 cytokines and 3 matrix metalloproteinases (MMPs), recognized for their contributions to the development of pulmonary fibrosis (**Figure S4**). Once again, dose-dependent analysis was employed to ensure that indicated cytokines or proteinases were definitively responding to the exposure compound, bleomycin, and to ensure that false positive responses were limited. An immunoassay revealed that 8 out of these 17 cytokine-proteinases were secreted in a differential and dose-dependent manner in response to bleomycin exposure (**Figure 4A, Figure S4**).

**Figure 4:**
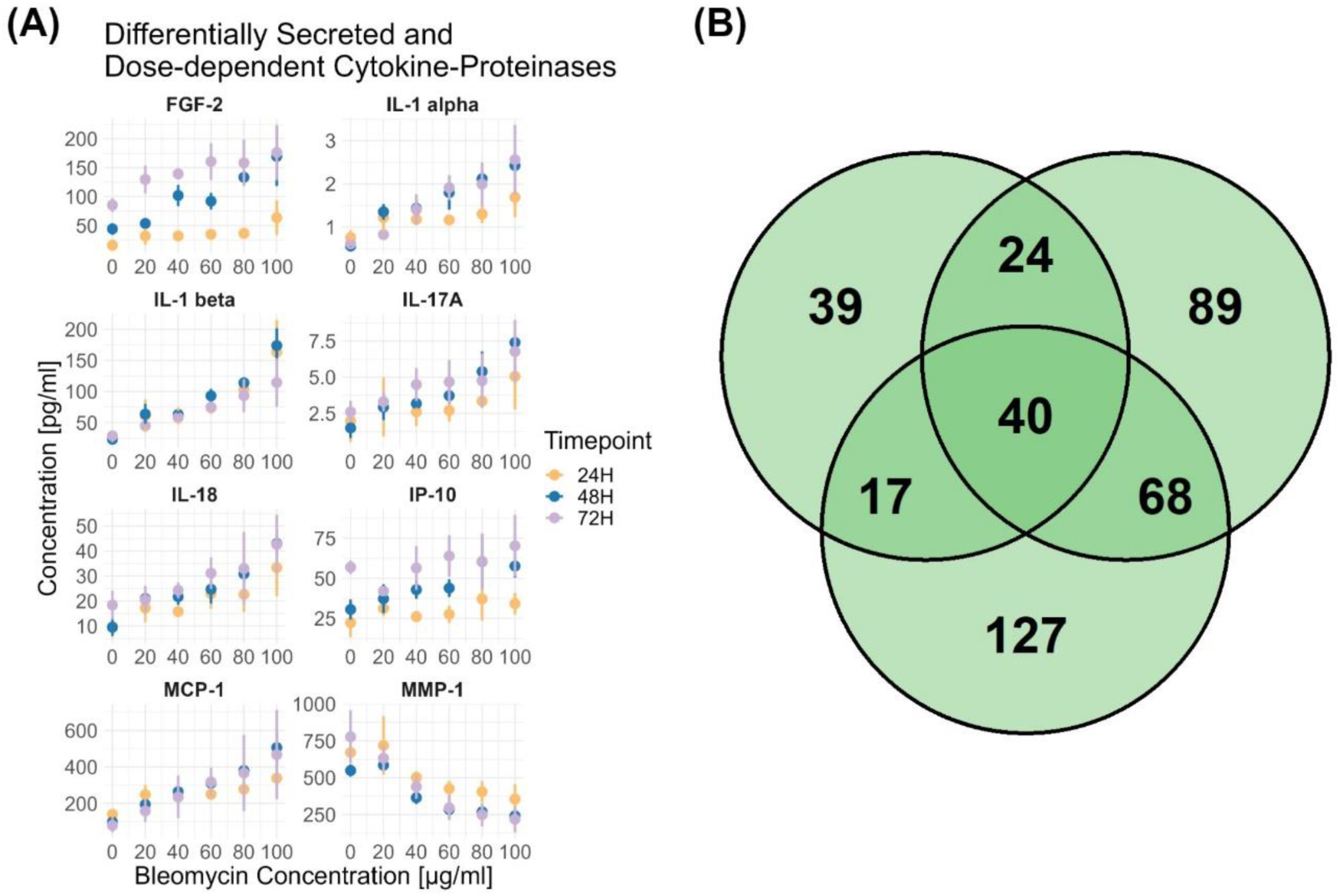
The correlation of differentially expressed (DE) and dose-dependent (DD) genes to a panel of cytokine-proteinases in THP-1 macrophages exposed to 0-100 µg/mL of Bleomycin at 24 H, 48 H and 72 H. (A) From a total of 17 cytokine-proteinases analysed, 8 were found to be differentially secreted (ANOVA padj <0.05, see Figure S4 and Supplementary File S1) and dose-dependent (see Figure S4). Scatter plots indicate the mean concentration of each protein (y-axis) across bleomycin concentrations (x-axis) for each timepoint. Whiskers represent standard deviation of around the mean (n=3-4). (B) Venn diagram shows differentially expressed (DE) and dose-dependent (DD) immune genes responding to bleomycin and correlating to cytokine-proteinase secretion at each timepoint (Spearman correlation −0.8<ρ>0.8, FDR adjusted p-value <0.05, see Supplementary File S1) filtered for immune genes in GO:0002376.

A dose-dependent increase in Fibroblast growth factor 2 (FGF-2) was seen at all 3 timepoints of bleomycin exposure, with the biggest increases at 48 H and 72 H at the highest concentration of bleomycin (**Figure 4A, Figure S4**). As well as having an obvious role in PF by acting as a potent mitogen in the differentiation and proliferation of myofibroblasts, FGF2 is also a strong activator of angiogenesis in respiratory disorders, and along with VEGF represents a potential treatment target in PF (66). Here, in our model of early-stage macrophage activity in PF, we also saw a dose-dependent increase in macrophage secreted IL-1 alpha, IL-1 beta, IL-17A and IL-18, sustained across 72 H of bleomycin exposure (**Figure 4A**). IL-1 beta, IL-17A and IL-18 are known to be increased in the bronchoalveolar lavage fluid of patients with IPF (21,22). We saw an increase in Interferon gamma-induced protein 10 (IP-10) in response to bleomycin exposure (**Figure 4A**). IP-10 is involved in the recruitment of monocytes during pulmonary inflammation events which can consequently advance fibrosis progression (67). Another monocyte recruiting signal is Monocyte Chemoattractant Protein-1 (MCP-1). The linear increase seen here in MCP-1 (also known as CCL2) with bleomycin exposure, is a signal by the macrophages to recruit and aggregate increased numbers of monocyte derived macrophages, to resolve epithelial damage (**Figure 4A**). The ablation of the MCP-1 receptor CCR2 in mice is known to limit the progression of bleomycin induced lung fibrosis by inhibiting the infiltration of monocytes into the lung (17,18).

Finally, out of the three MMPs measured (MMP-1, MMP-7, MMP-9, **Figure S4**), only MMP-1 (a.k.a Interstitial Collagenase) showed a dose dependent decrease in secretion (**Figure 4A, Figure S4**). It has previously been considered a paradox that in IPF patients, an elevated level of MMP-1 can be observed, and yet it is ineffective in degrading fibrillar collagens and contributing to the resolution of fibrotic foci in PF lungs (68,69). An advantage of the approach we take here, is that we are observing effects representative of the early, acute response to profibrotic insult. Our observation of decreased macrophage MMP-1 secretion caused by bleomycin (**Figure 4A**) illustrates an early response to profibrotic insult, whereas the increased MMP-1 seen in IPF patient lungs is a late stage adaptation that has come too delayed to effectively resolve the overaccumulation of ECM (68,69). In the early stages of lung fibrosis, the decrease in macrophage MMP-1 secretion we show here in response to profibrotic insult (**Figure 4A**), may well be contributing to fibrotic progression by decreasing the enzymatic degradation of collagen, allowing it to accumulate and contribute to ECM stiffness and scarring.

As cytokine and proteinase secretion is sustained by a transcriptional programme, we brought the analysis of these proteins into our framework of toxicological assessment of profibrotic substances by exploring the correlation between gene expression changes caused by bleomycin, and the secretion of these cytokine-proteinases. We performed a correlation analysis between all DE∩DD genes responding to bleomycin and the secretion levels of the 8 identified cytokine-proteinases also responding to bleomycin (**See Supplementary File S1**). The resultant correlating genes were filtered for genes included in GO:0002376, immune system process. Among the correlating immune-related genes, 40 were consistently DE∩DD across all timepoints (**Figure 4B, Table S1**), representing a significant subset of the 108 total DE∩DD immune genes identified in our study (**Figure 3B, Table S1**). This indicates that the DE∩DD immune genes we identified are closely linked to external signalling by macrophages in response to bleomycin exposure, reinforcing the likelihood that these genes are implicated in the acute response to profibrotic insult and are involved in the progression of PF. The close correlation we find between dose-dependent gene expression regulation and cytokine-proteinase secretion also shows the validity of our approach to chemical safety assessment to resolve the immunotoxic alterations caused by profibrotic substances.

### 3.4 A safety assessment framework utilising immunotoxic gene panels

Taken together, the panel of 108 DE∩DD immune genes showing sustained response to bleomycin (**Figure 3B**), 40 of which were correlated to known PF cytokine-proteinases that also responded to bleomycin (**Figure 4B**), indicate the immunotoxic propensity of bleomycin at the early stages of PF progression. There are 19 cytokines and cytokine receptors in our panel of 108, which, along with 22 immune genes representing signalling receptors and DE∩DD immune gene families are indicated in the heatmap in **Figure 5**. Each of these genes showed the same consistent direction of regulation (up or down regulation) over all 3 timepoints, suggesting a sustained regulation of these genes in FAMs. We performed qPCR validation for this subset of immune genes in a completely independent repeat of the bleomycin exposure experiment, that showed agreement with the RNA sequencing data, indicating consistency and reproducibility of our findings (**Figure 5**). Such reproducibility is an important element in the development of chemical safety assessment models.

**Figure 5:**
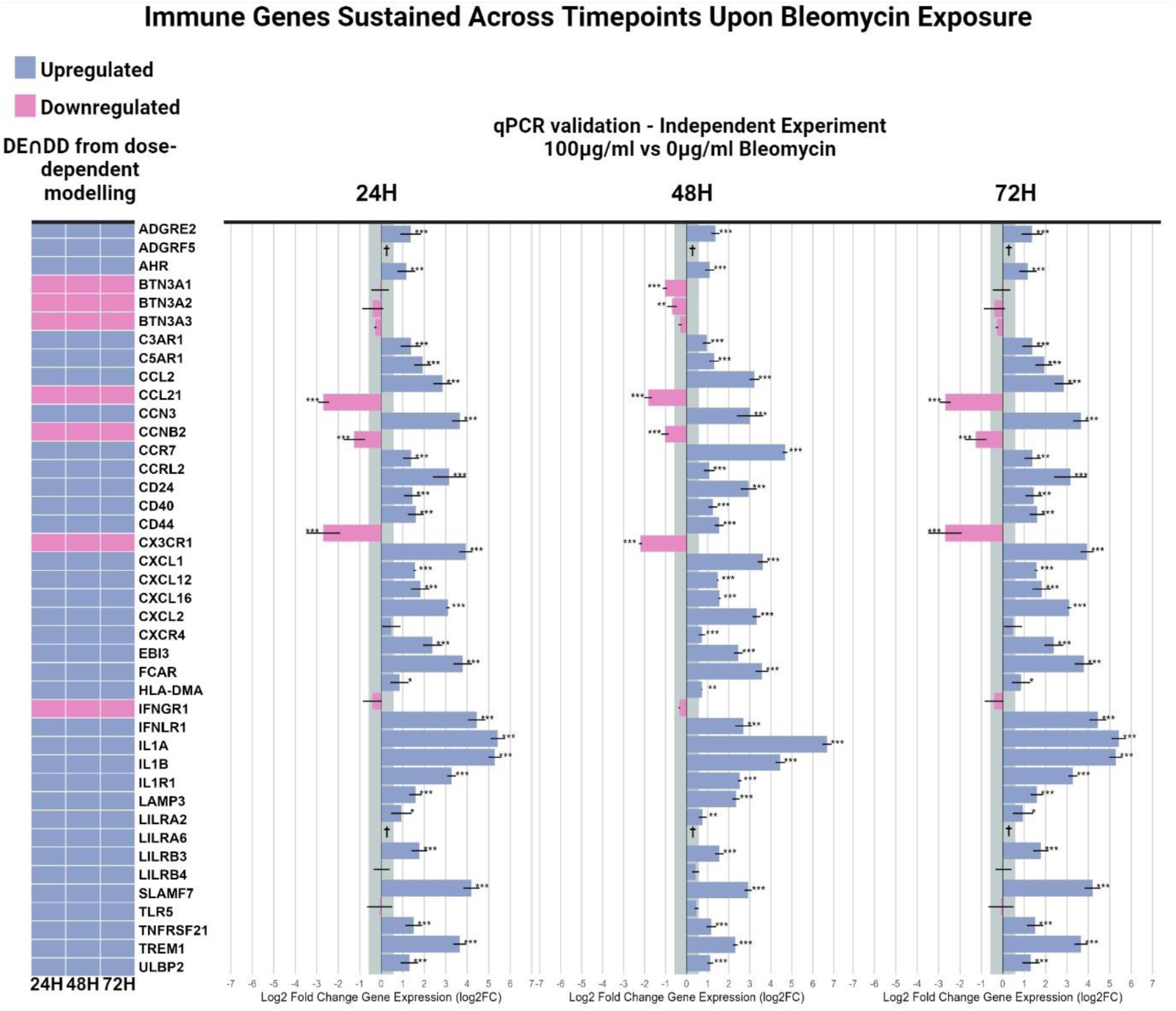
THP-1 macrophages exposed to 0-100 µg/mL of Bleomycin at 24 H, 48 H and 72 H have a distinct immune gene expression signature. A total of 108 immune genes were both differentially expressed and dose-dependently expressed across all three timepoints upon exposure to bleomycin. Of these genes the dose-dependent dysregulation direction (up or down regulated) of a select panel of cytokines and receptors, upon exposure to bleomycin is indicated in the heatmap across all 3 timepoints (24 H, 48 H, 72 H). An exact repeat independent bleomycin exposure of THP-1 macrophages was then performed and qPCR for these genes was performed, with results comparing the highest bleomycin concentration (100 µg/mL) to unexposed cells shown. Bars show Log2 Fold Change with std deviation. One-way ANOVA followed by Tukey’s HSD was performed (See Supplementary File S1). N = 4 in all cases. * p < 0.05, ** p < 0.01, *** p < 0.001. † = Gene was detected in 100 µg/ml bleomycin condition but undetected in 0 µg/ml bleomycin control condition preventing relative expression quantitation.

Whilst our panel of genes represents an immune signature in the very early exposure stages of PF, we also wanted to see whether these genes have been implicated in diagnosed PF patients representing the later stages of disease, as those immune genes that are dysregulated across all stages of fibrosis progression caused by an initial exposure to a profibrotic exposure, are likely to be of great importance. As such, we gave a score to each of the 108 DE∩DD immune genes based on whether they have been suggested to be implicated in pulmonary fibrosis in the DisGeNET database of genes and variants associated to human diseases, whether they are found in a set of GWAS data for IPF and whether they can be considered targets of currently approved drugs. We also took a set of 10 harmonised public RNA sequencing datasets representing biopsies from IPF patients vs healthy lung biopsies and assigned a score from 0 to 1 for each of the 108 DE∩DD immune genes in our panel based on how many times they were implicated in these biopsy experiments. Together with a score for if they were correlated to cytokine-proteinase secretion and if they were a cytokine or cytokine receptor themselves, we were able to rank the genes based on their importance in the immunotoxic signature of macrophages exposed to bleomycin (**Table S1**).

The top two highest ranked genes with the highest score were IL1B and C5AR1 (**Table S1**). C5AR1 and C3AR1 are G-protein coupled receptors that mediate innate immune complement activation pathways. Both receptors were dose-dependently up-regulated at all 3 timepoints of bleomycin exposure in FAMs (**Figure 5**). Antagonising these receptors has been shown to suppress bleomycin induced PF in mice by inhibiting TGFB levels and signalling activity (70,71). The known role of C5AR1 and IL1B in PF highlights the validity of our approach. The top-ranking genes in **Table S1** are therefore a representation of the immune activity of macrophages driving the pulmonary toxicity associated with bleomycin treatment that is sustained over the full course of progression to pulmonary fibrosis. Utilising the toxicological assessment framework established in this investigation, we were able to establish the immunotoxic profile induced by bleomycin in monocyte-derived macrophages. Whether this immunotoxic signature is also characteristic of other profibrotic agents remains to be elucidated. Nevertheless, the methodology applied herein offers a promising approach for fast and resource-efficient safety evaluation of other compounds suspected to induce fibrotic alterations.

Beyond using the methodology in this study as a safety assessment framework for profiling immunotoxic profibrotic potential, this study was also able to inform on the immune response to bleomycin. Targeting the 108 DE∩DD immune genes in our panel may well be a means to extend the use of bleomycin as an anti-cancer agent, or the expression of these genes might be explored for use during bleomycin treatment to monitor the likelihood of pulmonary side effects. When taken individually, these immune genes represent potential biomarkers for PF. An ideal biomarker should be measurable, reproducible, as well as specific and plausible for the disease or phenotype in question. On top of this, the biomarker should have a known temporal expression and a known gradient or dose-responsive expression to reflect the variability in underlying exposures or causes of the disease being looked at (72). Many of these properties apply to each of the panel of 108 DE∩DD immune genes. We looked at the baseline expression of these 108 DE∩DD immune genes in lung specific macrophages as compared to macrophages found in other tissues, using discriminant fuzzy pattern analysis on gene expression data for tissue specific macrophages, queried from the tabula sapiens dataset contained within a previously established knowledge graph **(Figure S5)** (53–55). When compared to other tissue macrophages, of the 74 out of 108 genes with expression data present, low baseline expression was seen in the majority of our panel of DE∩DD immune genes, where expression was often absent in macrophages from other tissues (**Figure S5**). This detectable but low baseline expression can also be considered an important attribute for useful biomarkers (72). Finally, the top ranked genes in **Table S1**, and their respective pathways, can also be considered potential targets for biologic treatments in PF due to their likely dysregulation at all stages of disease, but this requires further exploration beyond the scope of this study.

## 4. Conclusions

In this study, we exposed THP-1 macrophages to sublethal concentrations of bleomycin across multiple timepoints and analysed the resulting RNA sequencing data using a novel toxicogenomic approach that establishes dose-dependently expressed genes. By correlating the sequencing data with the expression profiles of a panel of cytokines known to be involved in pulmonary fibrosis, we were able to highlight immune genes involved in external immune signalling, modulated in response to a profibrotic stimulus. This methodology and safety assessment framework found a distinct immunotoxic profile, characteristic of macrophages subjected to bleomycin exposure. The relevance of this immunotoxic profile may extend beyond bleomycin exposure to encompass a broader range of profibrotic substances, and may even extend to other fibrotic diseases than PF. Therefore, the workflow of this study has the potential to be used for in vitro toxicity assessments for compounds with suspected fibrogenic effects. Our findings set the stage for further investigations to determine whether the immune signature we have identified is specific to bleomycin-induced fibrosis or if it holds broader applicability to other fibrotic triggers. Such research could solidify our approach as a standard framework for the development of novel safety assessment strategies for suspected profibrotic compounds.

## Author Contributions

Based on the CREDIT contributor roles taxonomy: **J.M.** contributed to conceptualization, formal analysis, investigation, methodology, project administration, supervision, validation, visualisation and writing – original draft. **A.F.** contributed to data curation, formal analysis and methodology. **L.M.** contributed to data curation and methodology. **S.I.** contributed to formal analysis. **A.P.** contributed to formal analysis. **S.S.** contributed to investigation and validation. **M.V.** contributed to investigation and resources. **S.P.** contributed to investigation. **A.S.** contributed to formal analysis, methodology and software. **D.G.** contributed to conceptualization, funding acquisition, project administration, resources, supervision, and writing – review & editing.

## Acknowledgements

This work received funding from the European Research Council (ERC) programme, Consolidator project ARCHIMEDES (grant agreement no. 101043848). J.M., A.F., and A.S. were supported by the Tampere Institute for Advanced Study.

In the preparation of this manuscript, ChatGPT-4.0 was utilized by J.M. for enhancing understandability and clarification of the text. All ideas and research are the original work of the authors.

## SUPPLEMENTARY FIGURES

**Figure S1:**
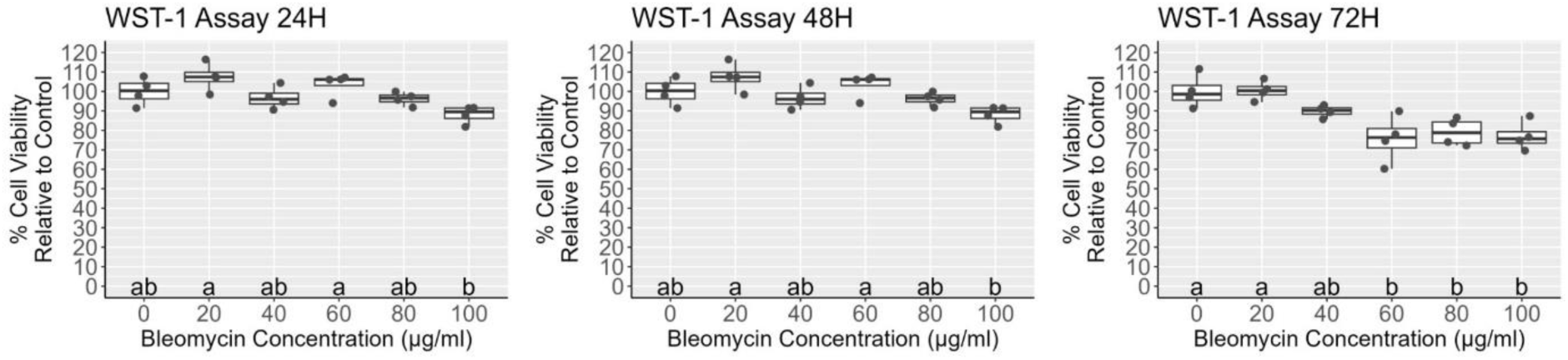
Viability of THP-1 cells exposed to 0-100 µg/mL of Bleomycin at 24 H, 48 H and 72 H. Points represent individual measures/wells of cells exposed to bleomycin with n = 4 for all concentrations and timepoints. Letters represent statistical categories (p < 0.05 between categories) from a tukey HSD posthoc test after One-Way ANOVA (Supplementary File S1). Boxplots show medians, with 25th and 75th percentile boxes, and minimum and maximum values as whiskers.

**Figure S2:**
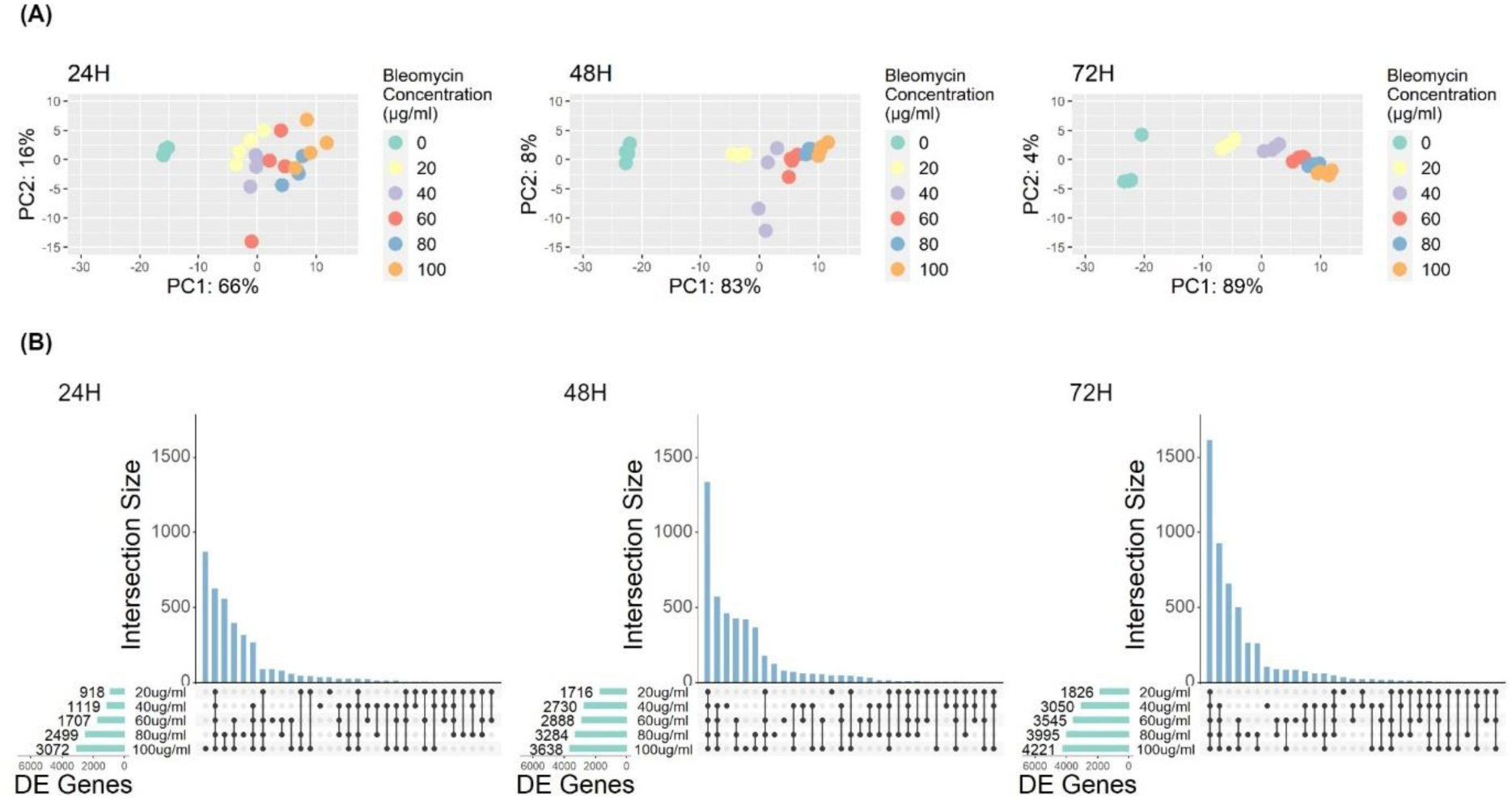
Differential Expression (DE,) of genes from THP-1 macrophages exposed to 0-100 µg/mL of Bleomycin at 24 H, 48 H and 72 H. (A) PCA plots of VST transformed counts (Deseq2 VST transformation) for all genes measured with RNA-seq. (B) Upset plots show the intersection of differentially expressed genes between concentrations (Blue vertical bars and Black intersect points) and show the number of genes differentially expressed at each concentration of bleomycin (Green horizontal bars) (Deseq2 differential analysis, FDR padj <=0.01, Log2FC = 0.58 (1.5FC)).

**Figure S3:**
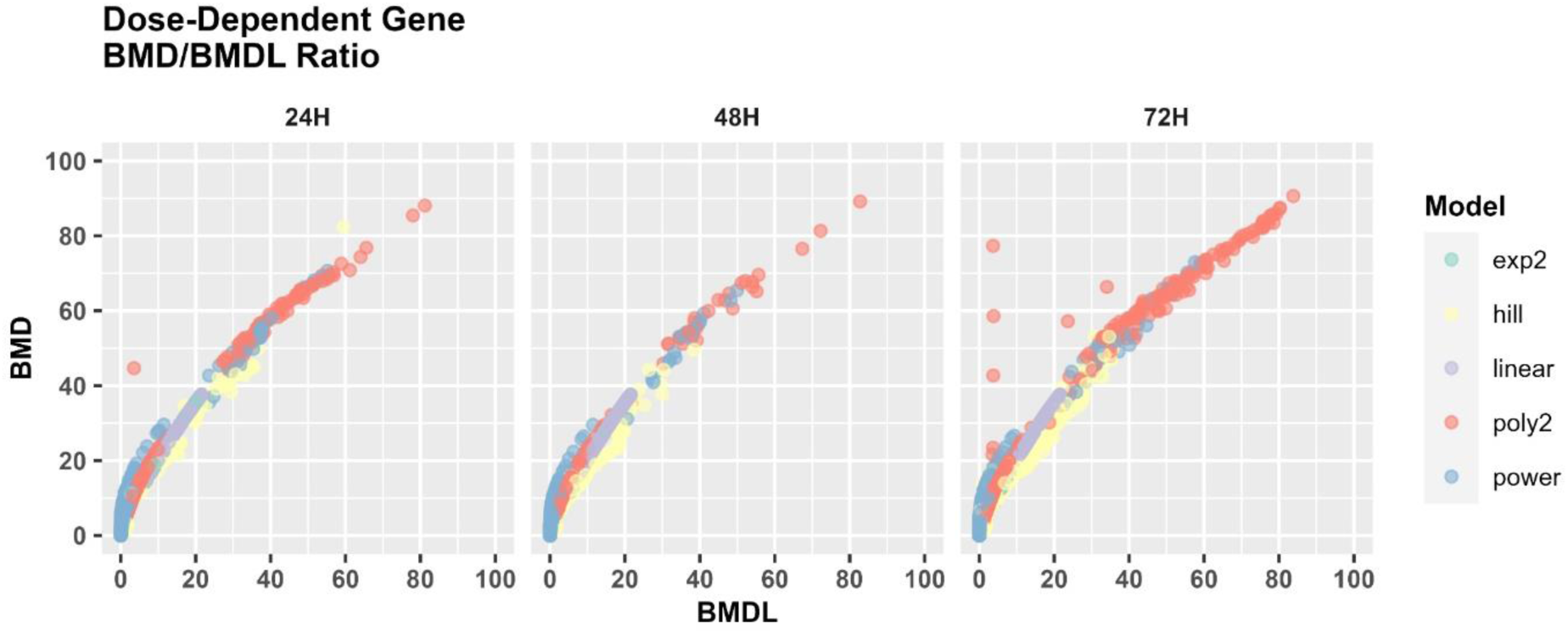
Dose-dependent (DD) analysis of genes from THP-1 cells exposed to 0-100 µg/mL of Bleomycin at 24 H, 48 H and 72 H. A scatter plot showing the ratio of benchmark dose (BMD) to benchmark dose lower confidence bound (BMDL) for each dose dependent gene with the model fitted to each gene indicated by colour.

**Figure S4:**
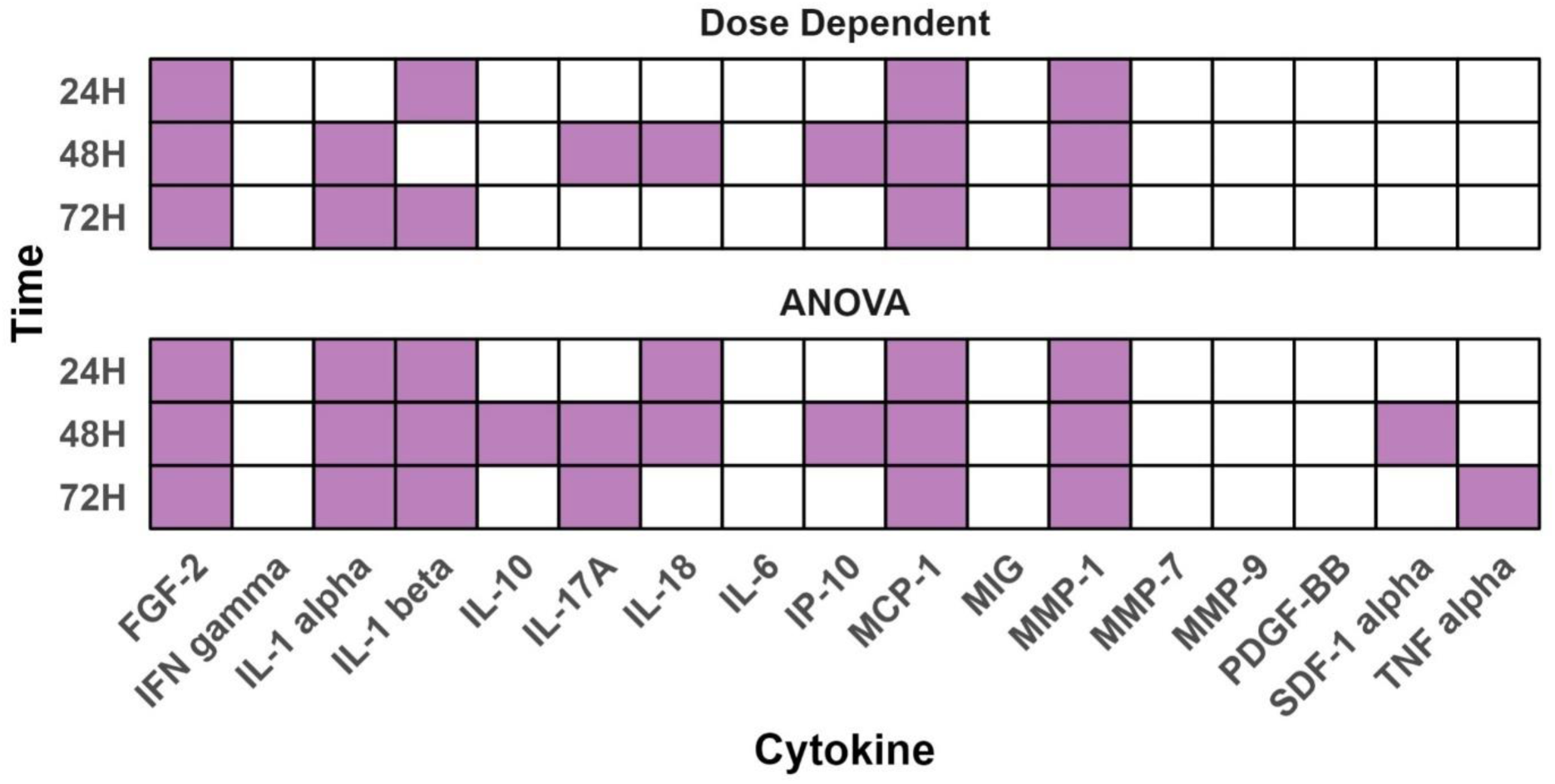
ANOVA and Dose-dependent analysis of a panel of cytokines and proteinases secreted from THP-1 macrophages exposed to 0-100 µg/mL of Bleomycin at 24 H, 48 H and 72 H. (A) From a total of 17 cytokines analysed, 11 were found to be differentially secreted (ANOVA padj <0.05, see Supplementary File S1) and of those, 8 were found to be dose-dependent, indicated by purple, at the timepoints shown.

**Figure S5:**
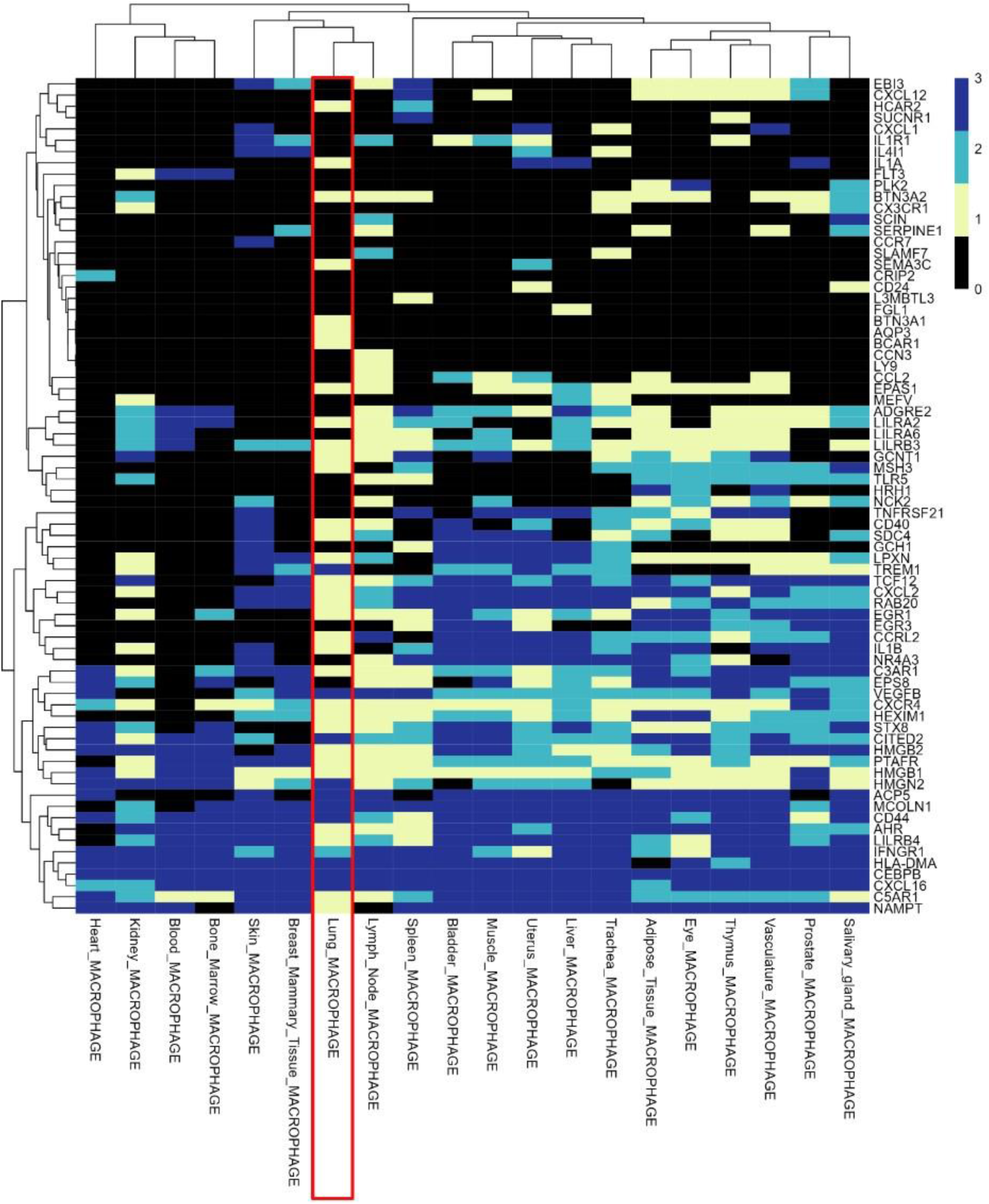
Heat map showing discriminant fuzzy pattern label expression values for 74 of 108 DE∩DD immune genes in macrophages from different tissues. 0 = no expression, 1 = low expression, 2 = medium expression, and 3 = high expression. Lung macrophages are highlighted with a red rectangle.

**TABLE S1:**
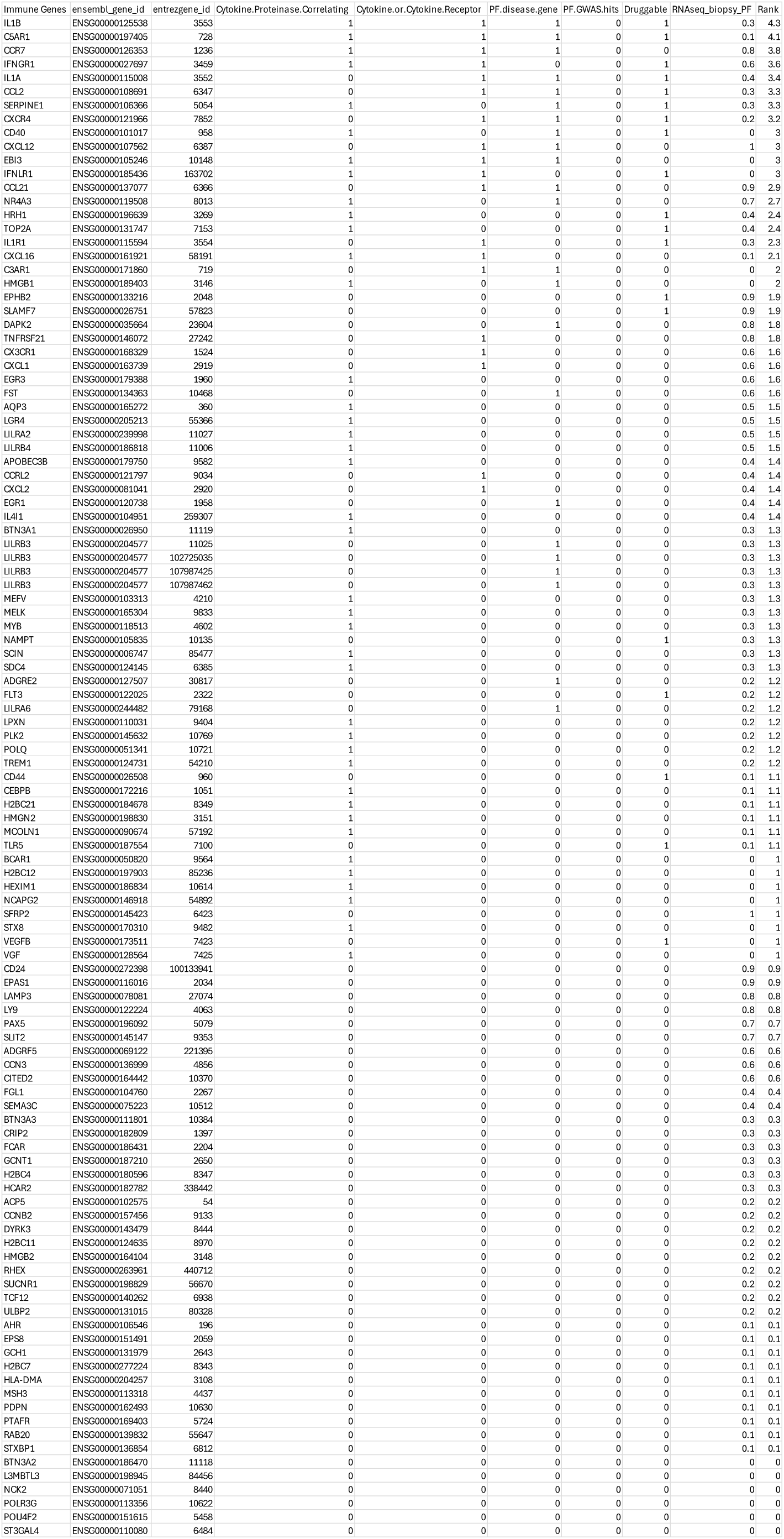
A ranked list of 108 differentially and dose-dependently expressed immune genes.

